# TIGIT-Fc Promote Immune Tolerance at the Feto-maternal Interface

**DOI:** 10.1101/819243

**Authors:** Wenyan Fu, Zetong Ma, Changhai Lei, Min Ding, Shi Hu

## Abstract

The perfect synchronization of maternal immune-endocrine mechanisms and those of the foetus is necessary for a successful pregnancy. In this report, decidual immune cells at the maternal-foetal interface were detected that expressed TIGIT (T cell immunoreceptor with Ig and ITIM domains), which is a co-inhibitory receptor that triggers immunological tolerance. We generated recombinant TIGIT-Fc fusion proteins by linking the extracellular domain of TIGIT and silent Fc fragments. The treatment with TIGIT-Fc of human decidual dendritic cells (dDCs) increased the production of interleukin 10 and induced the dDCs to powerfully polarize the decidual CD4^+^ T cells towards a classic T_H_2 phenotype. The administration of TIGIT-Fc to CBA/J pregnant mice at preimplantation induced CD4^+^ forkhead box P3^+^ (Foxp3^+^) regulatory T cells and tolerogenic dendritic cells and increased pregnancy rates in a mouse model of abortive stress. Moreover, we proposed that progesterone play a direct role in the transcriptional regulation of the TIGIT gene in decidual immune cell subsets. The results suggested the therapeutic potential of the TIGIT-Fc fusion protein in reinstating immune tolerance in failing pregnancies.

## Introduction

T cell immunoreceptor with Ig and ITIM domains (TIGIT, also known as Vstm3, VSIG9, and WUCAM) is a member of the immunoglobulin superfamily and belongs to the poliovirus receptor (PVR)/nectin family. Structurally, the N terminus of TIGIT is an extracellular immunoglobulin variable-set (IgV) domain, which is followed by a transmembrane domain and an intracellular domain. The intracellular domain of TIGIT contains a canonical immunoreceptor tyrosine-based inhibitory motif (ITIM) and an immunoglobulin tyrosine tail (ITT) motif (1). TIGIT was first identified in a genomic search for genes that encoded potential inhibitory receptors, which were identified according to the presence of certain protein domain structures that were expressed specifically in T cells (1). TIGIT expression is strictly limited to lymphocytes and shows the highest expression in follicular helper CD4^+^ T cells, effector and regulatory CD4^+^ T cells, effector CD8^+^ T cells, and natural killer (NK) cells(1–6). PVR, also known as CD155, Necl5, and Tage4, was identified as a cognate receptor for TIGIT with high affinity. Despite their weaker affinities, PVRL3 and CD112 (also known as PVRL2/nectin 2) were also shown to bind to TIGIT(1). A ‘lock and key’ trans-interaction between the TIGIT IgV domain and cis-homodimers of PVR was mediated by the distinctive (V/I)(S/T)Q, AX6G, and T(F/Y)PX1G submotifs(1, 3, 7), which define the PVR/nectin family comprising TIGIT, CD226, CD96, CD112R, PVR, CD112, and CD113 (also known as PVRL3/nectin 3)(1, 7–10). Nectins and nectin-like proteins are a group of surface receptors that function through homophilic and heterophilic trans-interactions and consequently mediate cell–cell adhesion, cell polarization, tissue organization, and signal transduction (11, 12).

Recently, an increasing number of mechanisms underlying TIGIT immune suppression have been identified. TIGIT can not only inhibit natural killer (NK) cell effector function but also suppress their dendritic cell costimulatory ability. The former blocks initial target cell death and the release of cancer-related antigens, and the latter results in increases in anti-inflammatory cytokines such as IL-10 and reduced target cell antigen presentation.

TIGIT could also stimulate PVR signalling on other cells, such as tumour cells. Suppressed CD8^+^ T cell effector function or skewed CD4^+^ T cell polarization could be provoked by TIGIT, PVR-stimulated myeloid cells, and TIGIT^+^ regulatory T cells (Tregs), which can also inhibit CD8^+^ T cells and prevent the elimination of target cells(13).

A successful pregnancy is a unique type of immunological process in which the semiallogeneic paternal antigens carried by the foetus are accepted by the maternal immune system, allowing trophoblasts to invade. Meanwhile, the defence mechanisms against pathogens in the maternal immune system are preserved. However, the mechanisms regulating these unique immunological behaviours and maintaining the harmonious coexistence of maternal- and foetal-derived cells remain poorly understood(14). Given that the dysregulation of maternal–foetal immunity and deficient placentation have a notable relationship with pregnancy loss and pregnancy complications, such as intrauterine growth restriction (IUGR)(15), recurrent spontaneous abortion (RSA)(16), and pre-eclampsia (PE) (17), further studies to advance the diagnosis and prevention of these conditions are urgently needed.

Up to 5% of all women attempting to conceive are affected by RSA, which is defined as two or more miscarriages(18). During the past few decades, growing evidence has proven the inevitable role of a misdirected maternal immune response in RSA. Because of the disturbance of haematological and immunological homeostasis, both autoimmune diseases and alloimmune disorders can create a uterine microenvironment that is difficult for the embryo and invading conceptus-derived placental trophoblasts. Disappointingly, current immunotherapies for RSA, including the use of hormones, antithrombotic drugs, intralipids, intravenous immunoglobulin (IVIG), cytokine agonists or antagonists, and allogeneic lymphocytes, have not consistently yielded successful pregnancy outcomes (19).

Previously, we showed that the administration of the TIGIT-Fc fusion protein to NZB/W F1 mice decreased the production of anti-double-stranded DNA antibodies, alleviated proteinuria and prolonged survival compared with that in mice treated with control IgG(20). The TIGIT-Fc fusion protein showed an IgG-like stability that was similar to that of CTLA-4-Fc. Here, we found that TIGIT was also expressed by decidual immune cells at the maternal-foetal interface during early pregnancy. The TIGIT-Fc fusion protein with silent Fc fragments guided dDCs to strongly polarize decidual CD4^+^ T cells towards a classic T_H_2 phenotype. In a mouse model, we obtained new experimental evidence to support the administration of TIGIT-Fc to promote fetomaternal tolerance and demonstrated the therapeutic potential of TIGIT-Fc to restore immune tolerance in failing pregnancies. Moreover, our data also supported a potential role of progesterone/progesterone receptors in the regulation of TIGIT gene expression in decidual lymphocyte cells.

## Material and methods

### Primary human cell assays

PBMCs and decidual samples were isolated from the same patients who underwent induced abortion. Under a stereomicroscope, decidual samples were carefully separated from villi, homogenized, filtered through a 32-μm nylon mesh, and ultimately purified from decidual mononuclear cells (leukocytes) with the standard Ficoll-Hypaque method, as reported previously (21, 22). PBMCs were also obtained with the Ficoll-Hypaque method. All specimens were collected by using a protocol approved by the Second Military Medical University Review Board, and written informed consent was obtained from each donor. Cells were sorted into different subsets with the appropriate magnetic beads or by flow cytometry. Magnetic cell sorting using a CD1c (blood DC antigen-1) DC isolation kit (Miltenyi Biotec, Bergish, Glandbach, Germany) was used to further purify the decidual DCs from decidual mononuclear cells. A monocyte isolation kit (Miltenyi Biotec, Bergish, Glandbach, Germany) was used to isolate CD14^+^ monocytes with a purity of over 95%. As described previously (23), immature DCs were obtained from cells incubated with IL-4 (R&D Systems) and recombinant granulocyte-macrophage colony-stimulating factor (R&D Systems). On day 5, a group of immature DCs with a specific phenotype (CD14^−^, major histocompatibility complex class II-positive, CD80^+^, CD86^+^ and CD83 l) comprised over 90% of the cells (clones M5E2, TU36, L307.4, IT2.2 and HB15e, respectively; BD Biosciences). Mature DCs (MDDCs) were treated with LPS (100 ng/ml; Sigma), TNF-α (100 ng/ml; Invitrogen), CD40L (100 ng/ml; R&D Systems), Pam3CSK4 (N-palmitoyl-S-[2,3-bis(palmitoyloxy)-(2RS)-propyl]-[R]-Cys-[S]-Serl-[S]-Lys4 trihydrochloride; 100 ng/ml; Invitrogen) or fusion proteins. Cell culture supernatants were collected to measure the cytokine concentrations with enzyme-linked immunosorbent assay (ELISA) kits specific for human IL-12p40, IL-12p70 and IL-10 (BD Bioscience and R&D Systems).

### Real-time polymerase chain reaction (PCR)

Total RNA was isolated with the RNeasy Mini Kit (Qiagen) according to the manufacturer’s instructions. Real-time quantitative PCR was performed with an ABI PRISM 7900HT instrument with the commercially available TaqMan probes Hs00545087 and Mm03807522. Data were normalized to β-actin, which served as an endogenous control, and analysed using SDS v2.3 (Applied Biosystems).

### Fusion proteins

As previously reported (1, 20), a recombinant plasmid was constructed by fusing the Fc segment of human IgG1 or murine IG2a, encoding the hinge-CH2-CH3 segment, to the C-termini of the extracellular domains (ECDs) of human and murine TIGIT, respectively. The LALA-PG Fc variant was constructed as previously described (24). All fusion proteins were obtained via the FreeStyle 293 expression system (Invitrogen) according to previously reported methods (25, 26) and subsequently purified using protein A-sepharose from the harvested cell culture supernatant. The purity of the fusion protein was determined by polyacrylamide gel electrophoresis. The protein concentration was measured according to the UV absorbance at a wavelength of 280 nm.

### Affinity measurement

By using standard amine-coupling chemistry, we immobilized an anti-murine Fc polyclonal antibody (Jackson ImmunoResearch Europe Ltd.) on a CM5 chip (~150 RU). Different antigens or antibodies (12.5 nM~200 nM) were injected to capture the fusion proteins by using a previously reported method (20). The measurement of the monovalent binding affinity of the fusion protein was calculated by using surface plasmon resonance (SPR) (BIAcore-2000). The kinetic analysis was performed using a 1:1 Langmuir model that simultaneously fit k_a_ and k_b_.

### IgG Biological Effect Assays

For the in vitro ADCC assay, SupT1 cells expressing murine PVR and A431 cells (high-expressing PVR cells) were labelled with 5 mM carboxyfluorescein succinimidyl ester (CellTrace CFSE Cell Proliferation Kit, Life Technologies) and co-cultured with murine or human macrophages overnight, respectively, at the indicated ratios in the presence of TIGIT-Fc fusion proteins.

For C1q ELISA, an ELISA sandwich-type immunoassay was used to analyse the binding of the different fusion proteins to C1q. Each fusion protein was coupled to a hydrophobic Maxisorp 96-well plate at eight different concentrations between 10 and 50 μg/ml. After washing, the C1q samples were incubated on the plate to allow C1q to bind to the fusion proteins. The bound C1q molecules were further washed and detected by anti-C1q antibodies followed by an HRP-labelled secondary antibody.

### Animal studies

C.B-17SCID;DBA/2J and CBA/J mice were provided by the Animal Center of the Second Military Medical University. All animals were treated in accordance with the guidelines of the Committee on Animals of the Second Military Medical University. The pharmacokinetic parameters (PK) of the fusion proteins were determined in female C.B-17 SCID mouse models. The fusion proteins were administered to eight-week-old mice at a dose of 1 mg/kg body weight by tail vein injection. Blood was collected in heparin-containing tubes and centrifuged to obtain the plasma samples. The serum concentration of the fusion proteins was determined by ELISA.

For the drug treatment studies, all mice were used at 10–12 weeks of age. To explore the protective role of TIGIT-Fc during pregnancy, an immunological model of abortion was used in which DBA/2J-mated CBA/J female mice were exposed to sound stress as previously described. Briefly, DBA/2J-mated CBA/J females were randomized and divided into three groups: group 1, CBA/J + control IgG; group 2, CBA/J + stress + control IgG; and group 3, CBA/J + stress +TIGIT-Fc. CBA/J mice were exposed to sound stress. Selected mice were treated with 10 mg/kg fusion proteins (i.v.) on 1.5 and 3.5 day d postcoitum(dpc). Paraaortic lymph nodes (PALN) and uterus cells were analyzed at 6.5 dpc. Thereafter, 8 mice per group were killed at 12 dpc to assess the pregnancy and abortion rates.

### Luciferase Reporter Assay and Chromatin Immunoprecipitation (ChIP)

To determine the effect of progesterone on TIGIT expression, cells were transfected with an expression vector encoding a full-length progesterone receptor and plasmids encoding a TIGIT-dependent luciferase reporter construct and a transfection control reporter (Renilla luciferase). The transfected cells were treated with progesterone or the antiprogestin mifepristone (10^−6^ M) in 0.01% (v/v) ethanol when indicated. The amount of luciferase was measured 48 hours later using a dual luciferase assay kit (Promega), and the firefly luciferase activity was normalized to the Renilla luciferase activity.

ChIP assays were performed as described previously (Dong et al., 2012; Lin et al., 2010). The primer sets used for the TIGIT and Tight promoters are listed in the supplemental Table 2.

### Statistical analysis

Unless otherwise specified, Student’s t test was used to evaluate the significance of the differences between two groups, and ANOVA was used to evaluate differences among three or more groups. Differences between samples were considered statistically significant when P < 0.05.

## Results

### TIGIT is expressed on decidual lymphocyte cells

A previous report showed that TIGIT was expressed in NK and T cells(1). We first detected TIGIT expression by quantitative RT-PCR in decidual immune cell subsets from early pregnancy (Fig. 1A). The high expression of TIGIT was detected in CD4^+^CD25^hi^ Treg cells, memory CD45RO^+^ cells and dNK cells, while naive CD45RA^+^ T cells, DC cells and decidual CD14^+^ monocytes/macrophages showed low levels of TIGIT mRNA expression. Further flow cytometry analysis confirmed that TIGIT expression was absent in decidual CD45RA^+^ CD4^+^ T cells and CD1c^+^ DC cells and was highest in CD4^+^CD25^hi^ Treg cells and CD45RO^+^ T cells (Fig. 1B).

**Figure 1.**
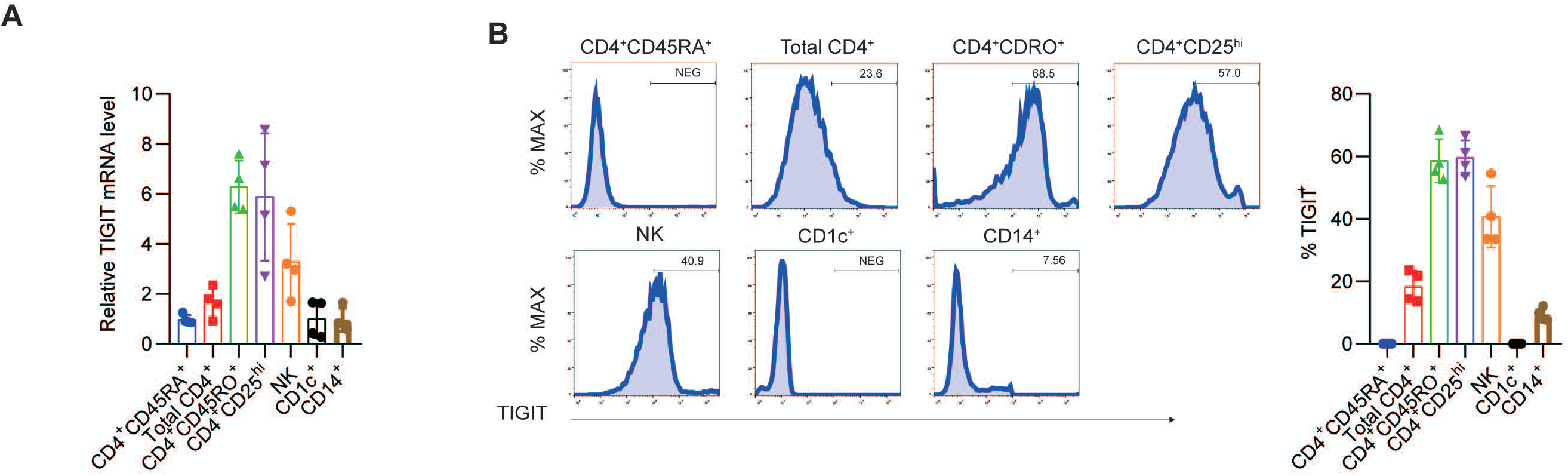
Expression of TIGIT protein in decidual immune cells. **(A)** qPCR of the expression of TIGIT mRNA in total CD4^+^, CD4^+^CD45RO^+^, CD4^+^CD45RA^+^, CD4^+^CD25hi (Treg), NK CD1c^+^ (DC) cells and CD14^+^ monocyte/macrophages relative to TIGIT expression in naive CD4^+^CD45RA^+^ cells. (B) Membrane-bound TIGIT expression in different subsets of immune cells. The expression of TIGIT on different human immune cells was detected by staining with the indicated antibody, followed by flow cytometry analysis. The proportions of cells with positive TIGIT expression are shown on the right (n = 4 independent biological experiments).

### TIGIT fusion proteins with silent effector Fcs

To investigate the therapeutic potential of TIGIT, we developed and generated a fusion protein by linking the extracellular domain of human or murine TIGIT to the human IgG1 Fc region (hTIGIT-Fc_wt) or murine IgG2a Fc chain (mTIGIT-Fc_wt), respectively. The antibody Fc region regulates the antibody serum half-life and cytotoxic activity. However, the TIGIT functional receptor, PVR, is ubiquitously expressed in the human placenta (27). Within the relevant therapeutic context, the cytotoxicity of an antibody is not desirable and can lead to safety issues by initiating native host immune defences against cells with receptor antigen expression. Therefore, we used LALA-PG Fc variants (hTIGIT-Fc_ LALA-PG; mTIGIT-Fc_ LALA-PG) that block complement binding and fixation as well as Fc-γ-dependent, antibody-dependent, and cell-mediated cytotoxity caused by both murine IgG2a and human IgG1. As previously reported, the fusion proteins showed high affinity for binding to CD155 (Fig. S1). We also found that hTIGIT-Fc bound to murine CD155, but such binding could not be detected between mTIGIT-Fc and human CD155 (Fig. S1). We next assessed the capacity of these fusion proteins to deplete PVR-expressing cells co-cultured with monocyte-derived macrophages in vitro at various effector to target (E:T) cell ratios (Figure 2A and B). As predicted, the hTIGIT-Fc_wt or mTIGIT-Fc_wt fusion proteins demonstrated strong ADCC activity, while the Fc-silent LALA-PG proteins showed negligible effects. In the C1q binding assays, only fusion proteins with the wild-type Fc showed remarkable binding to the C1q protein, which is part of the complement cascade (Fig. 2C and D). All LALA-PG protein variants were devoid of any detectable binding at protein concentrations of up to 50 μg/ml. In comparison with the Fc fusion protein CTLA4-Fc, which has been well studied in previous studies, the TIGIT-Fc fusion proteins exhibited IgG-like stability and a similar denaturation temperature. The lowest concentrations (< 2%) of low molecular weight and high molecular weight products were observed after storage at 1 mg/mL at 40 °C for 3 weeks (Table S1). A single intravenous dose of TIGIT-Fc proteins and CTLA4-Fc were separately administered to mice to measure the pharmacokinetic (PK) parameters. The main PK parameters of TIGIT-Fc proteins and CTLA4-Fc were very similar in mice and indicated the advantageously high stability of the TIGIT-Fc fusion proteins (Table S1). These data show that the hTIGIT-Fc_ LALA-PG and mTIGIT-Fc_ LALA-PG Fc variants do not induce any FcγR- or complement-mediated effector functions. Therefore, we used these proteins in the following experiments.

**Figure 2.**
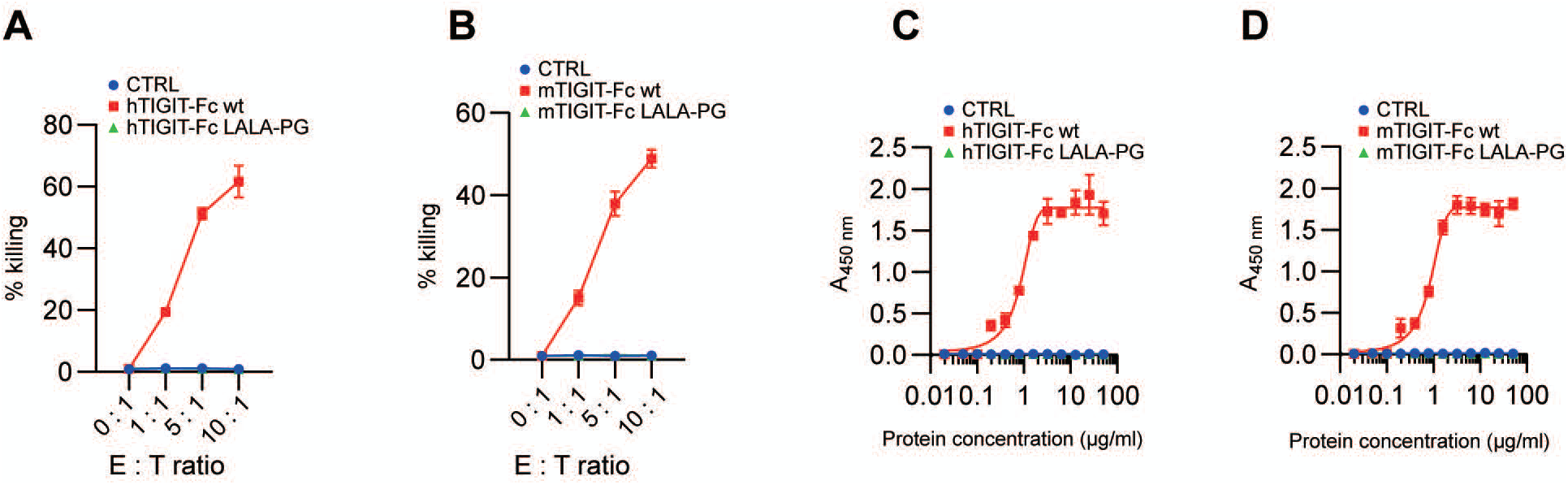
Biological effect of TIGIT fusion proteins in vitro. **(A and B)** In vitro ADCC assay with monocyte-derived macrophages and PVR^+^ target cells in the presence or absence of human (A) or murine (B) TIGIT-Fc fusion proteins. **(C and D)** C1q binding assay. Human (C) and murine (D) TIGIT-Fc fusion proteins were coated on an ELISA plate. The complement component C1q was added to the plate, and its retention was detected via anti-C1q antibodies.

### TIGIT-Fc modifies decidual DC cytokine production

Treatment with TIGIT-Fc during DC maturation influenced DC cytokine production (1). We tested whether TIGIT-Fc with a silent IgG has a similar effect on cytokine production. Our data showed that in monocyte-derived DCs (MDDCs) stimulated with TNFα, TIGIT-Fc significantly enhanced the production of IL-10 and inhibited the production of IL-12p40 and IL-12p70 (Fig. 3A). IL-12 production by MDDCs matured by CD40L or the TLR4 ligand lipopolysaccharide (LPS) was also significantly decreased. These data are consistent with those in a previous report. We next tested the effects of TIGIT-Fc on dDCs during maturation. Interestingly, in dDCs stimulated with TNF-α, CD40L or LPS, TIGIT-Fc also significantly increased the production of IL-10 and inhibited the production of IL-12 (Fig. 3B). Moreover, the cytokine production of MDDCs matured by the TLR2 ligand Pam_3_CSK_4_ was not significantly affected by TIGIT-Fc. The maturation status of MDDCs or dDCs, as defined by the surface expression of CD80, CD86, CD83 and HLA-DR, was not affected by TIGIT-Fc (Fig. 3 C and D).

**Figure 3.**
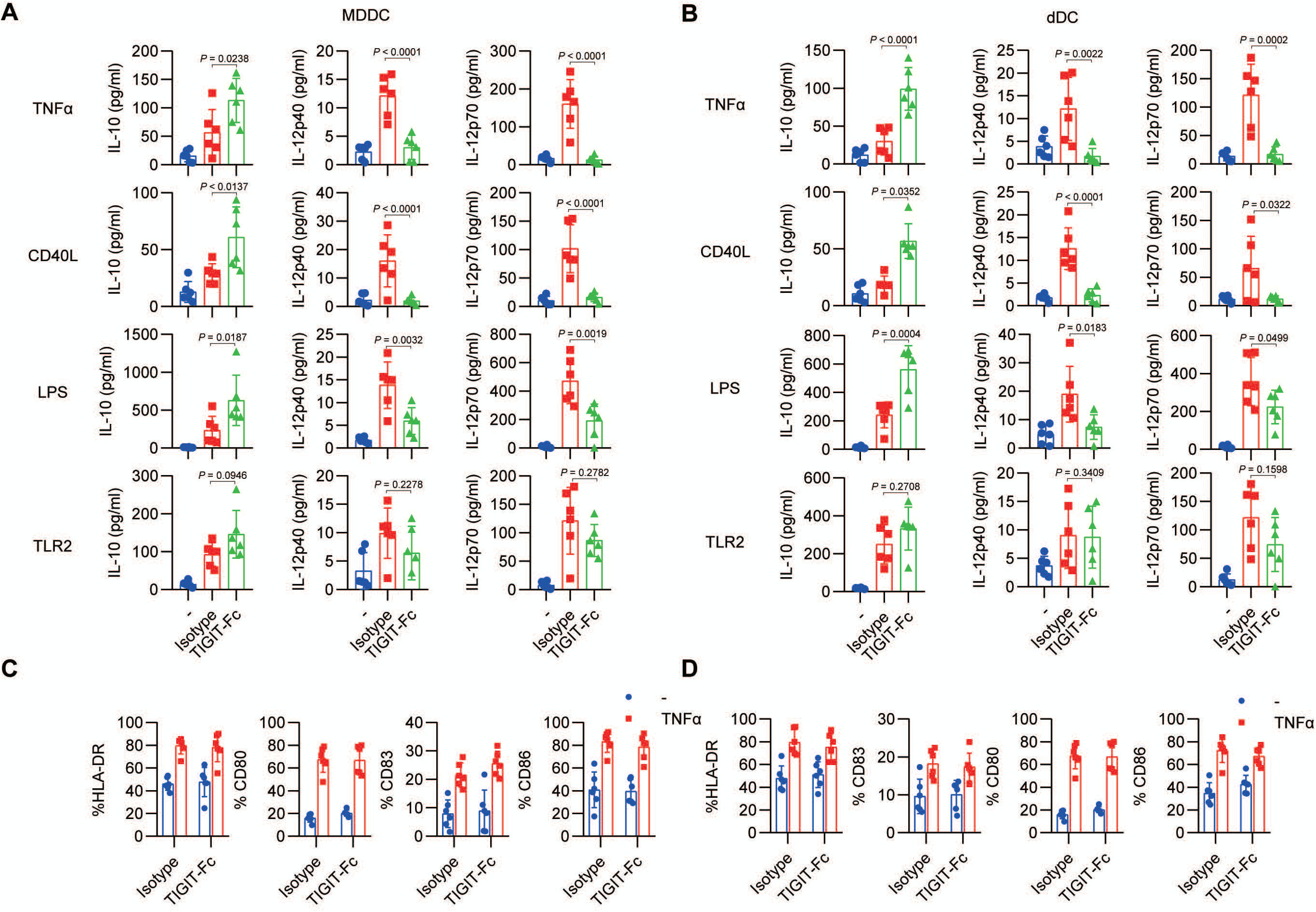
Modification of DCs by TIGIT-Fc during maturation. **(A and B)** ELISA of cytokine production in 2 × 10^5^ immature MDDCs (iMDDCs) (A) or dDCs (B) matured for 24 hours with TNF-α, CD40L, LPS or Pam3CKS4 (TLR2 ligand) or cultured in medium alone (iMDDCs or dDCs;-) in the presence of hTIGIT-Fc or an IgG1-variant isotype-matched control antibody (10 μg/ml); **(C and D)** Flow cytometry of cell surface markers on iMDDCs (C) and dDCs (D) incubated for 24 hours in medium alone (–) or with TNF-α in the presence of hTIGIT-Fc or an isotype-matched control antibody, represented as the proportion of cells with positive markers. Data are from one of six independent experiments. Error bars, s.d.

### TIGIT induced decidual DCs to secrete high levels of IL-10 and polarized decidual CD4^+^ T cells towards a T_H_2 phenotype

Decidual DCs were reported to play a crucial role in tolerance at the maternal/foetal interface and thus played a pivotal role in preventing the immune rejection of the foetus. Next, we investigated whether TIGIT-Fc alone could induce decidual DCs to produce high levels of IL-10. Freshly isolated CD1c^+^ decidual DCs and peripheral blood MDDCs were treated with TIGIT-Fc (10 μg/mL) or LPS (100 ng/mL) for 48 hours. Our data show that that IL-10 but not TNF-α secretion by decidual DCs was strongly stimulated by TIGIT-Fc (Fig. 4A). An increase in IL-10 secreted by MDDCs from early pregnancy was not detected, suggesting a functional difference between MDDCs and dDCs.

**Figure 4.**
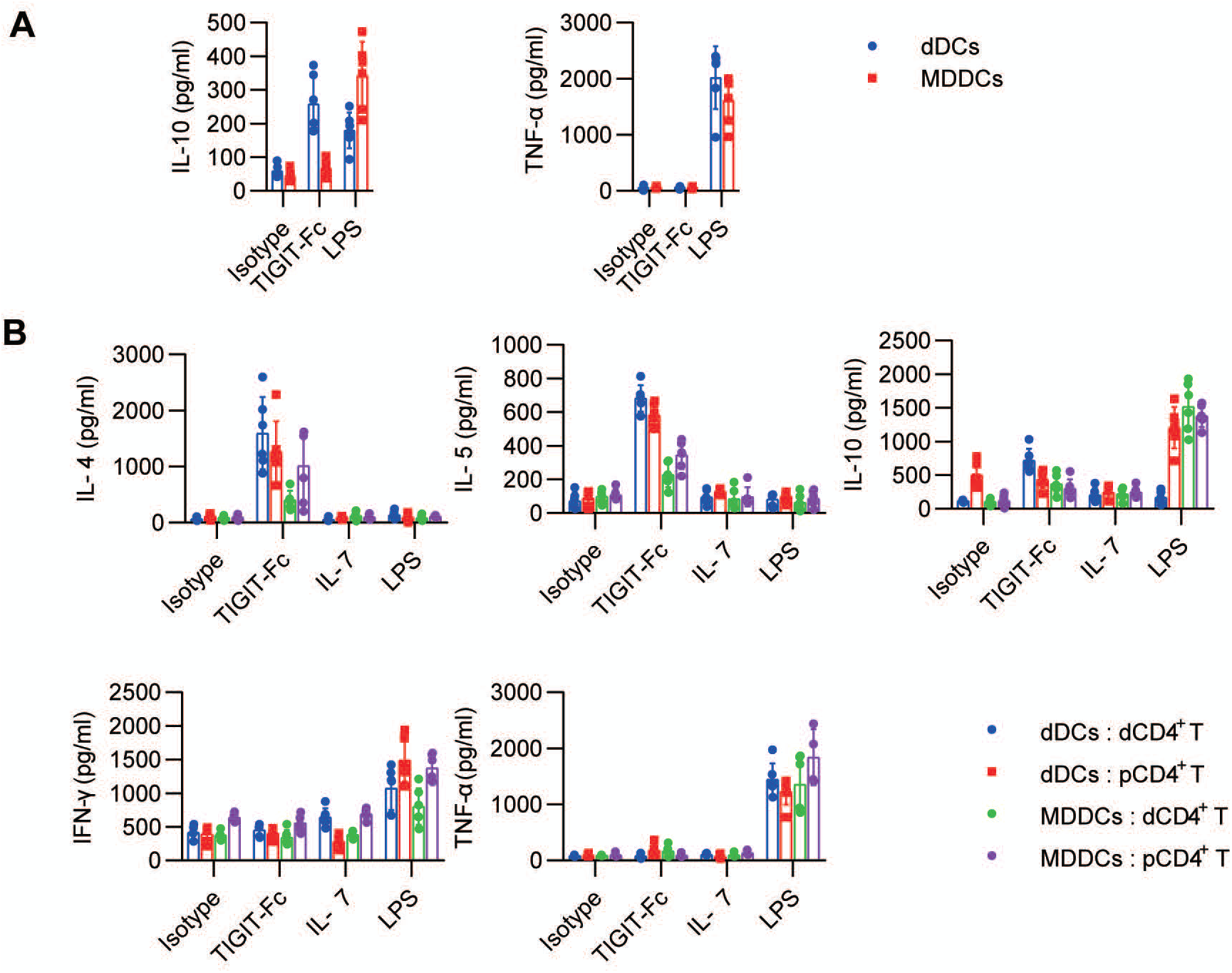
TIGIT-Fc administration causes DCs to induce CD4^+^ T cells to polarize towards a T_H_2 phenotype. **(A)** dDCs or MDDCs were treated with hTIGIT-Fc or LPS for 48 hours. The expression levels of IL-10 and TNF-α were measured by ELISA. **(B)** The dDCs and MDDCs were treated with TIGIT-Fc, IL-7, or LPS, respectively, for 24 hours, washed and cocultured with decidual CD4^+^ T (dCD4^+^ T) or peripheral CD4^+^ T (pCD4^+^ T) cells, respectively. The T cells were then stimulated, and the cytokine secretion of the CD4^+^ T cells was determined by ELISA. Data are the mean ± SD from 6 independently conducted experiments.

Next, we investigated whether TIGIT-Fc-induced dDCs could polarize decidual CD4^+^ T cells towards a T_H_2 phenotype. MDDCs or dDCs pretreated with TIGIT (10 μg/mL), IL-7 (100 ng/mL), or LPS (100 ng/mL) for 48 hours were cocultured with allogeneic peripheral or decidual CD4^+^ T cells for 3 days, respectively. IL-4, IL-5, IL-10, IFN-γ, and TNF-α in the supernatant were measured by ELISA (Fig. 4B). When dDCs were cocultured with decidual CD4^+^ T cells, IL-4 IL-5 and IL-10 were markedly upregulated by TIGIT-Fc treatment, while IFN-γ and TNF-α were not upregulated compared with that in decidual CD4^+^ T cells cocultured with dDCs but without TIGIT-Fc. However, we did not observe similar phenomena for IL-7- or LPS-activated dDCs, indicating that TIGIT-F-stimulated dDCs can polarize decidual CD4^+^ T cells towards a T_H_2-biased profile. The pretreatment of peripheral MDDCs from early pregnancy with TIGIT-Fc also promoted the high production of IL-4, IL-5, and IL-10 by peripheral CD4^+^ T cells, but the effect was not as strong as that observed for dDCs. These results indicated that treatment with TIGIT-Fc could influence the T_H_1/T_H_2 balance at the maternal-foetal interface.

### TIGIT Reduces Stress-Challenged Foetal Resorption by the Expansion of Foxp3^+^ Tregs

A well-established mouse model of stress-induced pregnancy failure that was reported previously (28) was used to explore the role of mTIGIT-Fc in fetomaternal tolerance. In the model, sound stress was used in DBA/2J-mated CBA/J females to challenge pregnancy maintenance and finally provoke foetal loss. Notably, a reduced incidence of foetal resorption (Fig. 5A) but an unchanged the number of foetal implantations (Fig. 5B) was observed in the stressed DBA/2J-mated CBA/J females administered mTIGIT-Fc. The phenotype of uterine CD11c^+^ cells was detected for further examination of the inhibitory effects of TIGIT-Fc on the stress-induced aberrant maturation of CD11c^+^ cells. Compared with the control group, pregnant DBA/2J-mated CBA/J females exposed to sonic stress were characterized by the maturation of uterine CD11c^+^ cells and showed increased levels of MHC-II and CD80. Treatment of the stress-challenged mice with mTIGIT-Fc decreased the number of phenotypically mature uterine CD11c^+^ cells (Fig. 5C). Additionally, the PALNs of mTIGIT-Fc-treated female mice showed a higher proportion of Foxp3^+^ cells at 6.5 dpc than that of control females (Fig. 5D). Overall, the results of our study indicate that TIGIT-Fc protects embryos from maternal immune rejection by inducing tolerant DCs (tDCs) and Foxp3^+^ cells.

**Figure 5.**
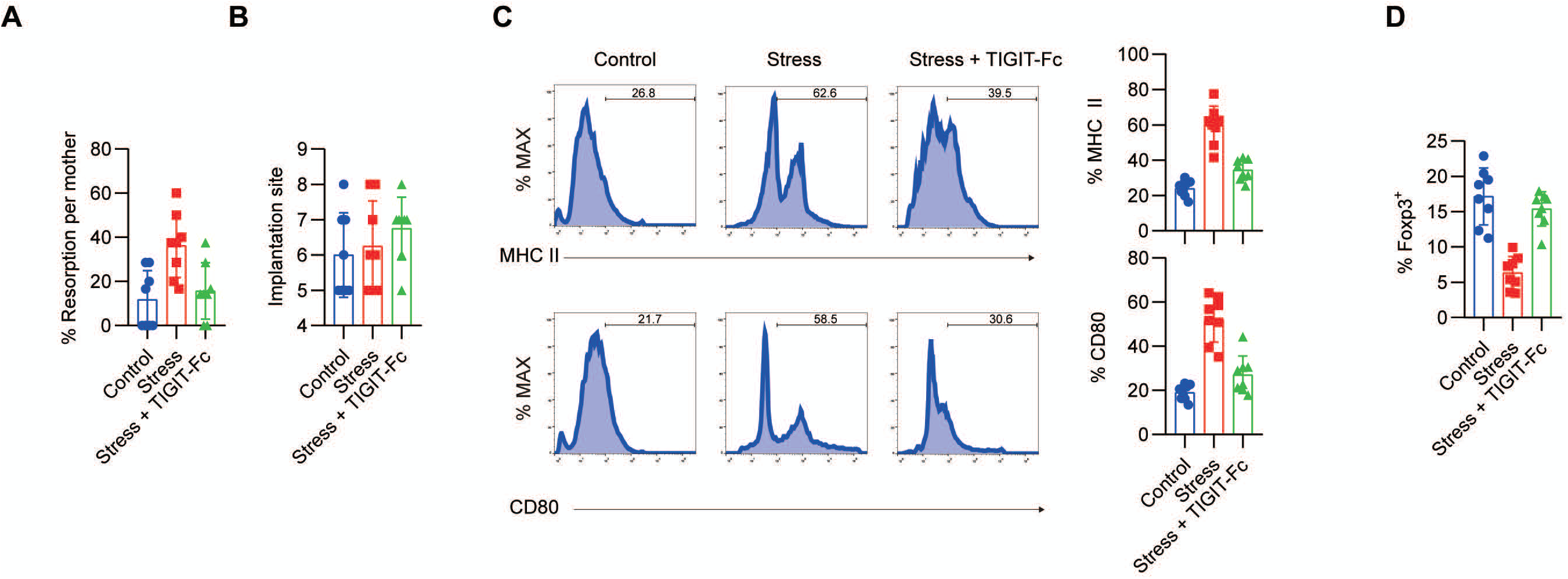
Administration of TIGIT-Fc protects foetuses from abortion. Treatment with mTIGIT-Fc of pregnant CBA/J females mated with DBA/2J males at 1.5 and 3.5 dpc. Sonic stress was applied from 2.5 dpc until sacrifice. **(A)** Resorption rates and **(B)** the total number of implantation sites were measured at 12.5 dpc in the mated female mice exposed to sound stress with or without mTIGIT-Fc. Bars, mean ± SEM. **(C)** Phenotypic analysis of CD11c^+^ uterine DCs from non-stressed mated mice treated with control IgG (Control), stressed mated mice treated with control IgG (Stress), and stressed mated mice treated with TIGIT-Fc (Stress + TIGIT-Fc) at 6.5 dpc. The percentages of MHC-II- and CD80-positive cells are plotted as bar graphs (right panels). Bars, mean ± SD. **(D)** Percentages of CD4^+^Foxp3^+^ T cells in the PALNs of different groups were calculated by flow cytometry analysis at 6.5 dpc.

### TIGIT expression is regulated by progesterone

To explore whether progesterone affects the transcription of TIGIT, the expression level of TIGIT mRNA in progesterone-treated CD4^+^ T cells and dNK cells was analysed with a qPCR assay. The results suggested that progesterone significantly upregulated *TIGIT* mRNA expression in a concentration-dependent manner. In comparison with the vehicle-treated control, progesterone at concentrations of 10^−7^ and 10^−6^ M increased the expression level of TIGIT mRNA in CD4^+^ T cells by 1.7- and 3.1-fold, respectively, and that in dNK cells by 1.8- and 4.7-fold, respectively (Fig. 5A). In a mouse model given the same treatment, the *Tigit* mRNA level in murine CD4^+^ T cells was increased by 1.2- and 2.3-fold, while in murine dNK cells it was increased by 2.5- and 4.3-fold, respectively, compared with that in the control group (Fig. 5B). The effect of progesterone (10^−6^) on the expression level of TIGIT and Tigit mRNA in both groups of cells was reversed by co-treatment with mifepristone (10^−5^M) (Fig. 6A, 2B), which verified that progesterone regulates TIGIT expression via progesterone receptors.

**Figure 6.**
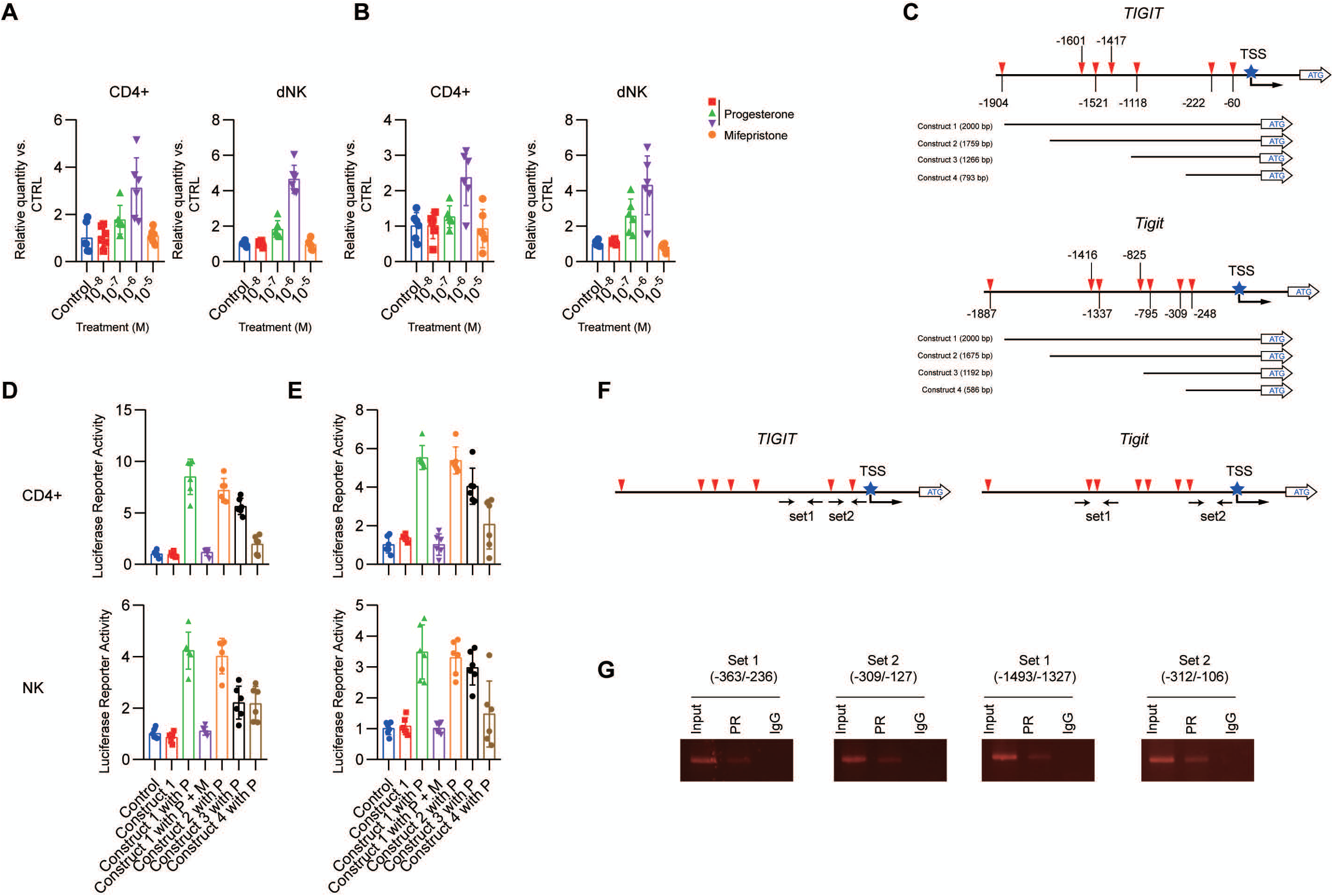
TIGIT is a target gene of progesterone. **(A and B)** qPCR of the expression of human *TIGIT* (A) or murine *Tight* (B) mRNA in CD4^+^ T and NK cells, which were treated with different concentrations of progesterone (10^−8^–10^−6^ M) or progesterone (10^−6^M) combined with mifepristone (10^−5^M) for 72 hours. (**C**) Schematic diagram showing the positions of the potential PREs in the TIGIT promoter (top) and *Tigit* promoter (bottom) in different promoter luciferase constructs. **(D)** *TIGIT* promoter luciferase constructs were expressed in CD4^+^ T cells and dNK cells, which were treated with progesterone or progesterone combined with mifepristone for 48 hours. Luciferase activities were determined and normalized. P, progesterone. M, mifepristone. **(E)** *Tigit* promoter-luciferase constructs were expressed in CD4^+^ T cells and dNK cells that were treated with progesterone or progesterone combined with mifepristone for 48 hours. Luciferase activities were determined as shown in (D). **(F)** Sets of primers used for the *TIGIT* and *Tight* promoter ChIP assays are shown. **(G)** The association of progesterone receptor and gene promoters in dNK cells treated with progesterone were analysed by ChIP.

Upon performing a computer-assisted homology search analysis, the result showed that TIGIT (−2000 to ^+^1) harbours seven half-consensus putative PREs upstream of the translation start site: PRE I (5’-TGTTCT-3’, located at −60 to −65); PRE II (5’-GGGACA-3’, located at −222 to −227); PRE III (5’-GGGACA-3’, located at −1118 to −1123), PRE IV (5’-GGGACA-3’, located at −1417 to −1422); PRE V (5’-TGTTCT-3’, located at −1521 to −1526); PRE VI (5’-TGTTCT-3’, located at −1601 to −1606); and PRE VII (5’-TGTTCT-3’, located at −1904). In Tigit, seven half-consensus PRE sites were also identified upstream from the translational start codon of Tigit: PREI (5’-AGAACT-3’, located at −248 to −253); PRE II (5’-TGTGCG-3’, located at −309 to −314); PRE III (5’-TGTGCA-3’, located at −795 to −800); PRE IV (5’-GGGACA-3’, located at −825 to −830); PRE V (5’-TGTTCT-3’, located at −1337 to −1342); PRE VI (5’-GGGACA-3’, located at −1416 to −1421); and PRE VII (5’-AGAACA-3’, located at −1887 to −1892).

We cloned both the human TIGIT and murine Tigit promoters (construct 1 = −2000 to ^+^1 bp) and developed several deletion mutants by using both constructs according to the location of these PREs (Fig. 6C). The significant upregulation of TIGIT promoter activity by progesterone was detected in CD4^+^ T cells and dNK cells with exogenous expression of the full-length TIGIT promoter reporter (construct 1) (Figure 6D), while this phenomenon was reversed by co-treatment with mifepristone. For the human TIGIT promoter reporters, when the PRE at −1904 was deleted (construct 1 versus construct 2), no change in TIGIT promoter-luciferase expression mediated by progesterone was observed, indicating that the PRE at −1904 is of little significance for the mediation by progesterone of TIGIT promoter-luciferase expression. However, four PREs upstream of the transcriptional starting site (TSS) (−1601, −1521, −1417 and −1118) were critical for progesterone-induced expression, as deletion constructs (construct 3 and construct 4) that removed these regions separately showed less sensitivity to progesterone-induced expression. For murine Tigit promoter reporters, our results showed that the PRE at −1887 was not critical for progesterone-mediated TIGIT promoter-luciferase expression (construct 1), while the deletion of the four PREs (−1416, −1337, −714 and −795) caused significant loss of progesterone-mediated luciferase activity, suggesting a crucial role for these sites (construct 4).

To verify whether the progesterone receptor binds to the TIGIT promoter, we performed chromatin immunoprecipitation (ChIP) by using two sets of primers for the human and murine gene promoters (Fig. 6F). Primer sets covering nucleotides from −630 to −449 and −450 to −288 in the human promoter as well as −1347 to −1114 and −241 to −14 in the murine promoter worked effectively and thus were used for the subsequent ChIP experiments. In dNK cells, we found that the endogenous progesterone receptor bound to the TIGIT promoter in human cells and murine cells (Fig. 6G). These results indicate that TIGIT is a direct target of the progesterone receptor.

## Discussion

As shown in previous studies, CTLA4-Ig, a fusion protein with the Fc segment of IgG1 linked with the extracellular domain of CTLA4, downregulated and blocked the CD28-B7.1/B7.2 co-stimulatory signal in T cells (29). Abatacept is a first-generation CTLA4-Ig co-stimulation blocker used for the treatment of rheumatoid arthritis(30). Compared to the control, the CTLA4-Ig-treated SLE mouse model had a delayed onset of SLE, the reduced production of autoantibodies, reduced proteinuria, and prolonged survival (31). The TIGIT/CD226-CD155-CD112 and CTLA-4/CD28-CD80-CD86 signalling pathways showed similar characteristics. The homologous molecules CD155 and CD112 prohibit the activation of T cells and NK cells by binding with the co-suppressor molecule TIGIT, reversely promoting activation by binding with the corresponding co-stimulatory molecule CD226. Likewise, the homologous molecules CD80 and CD86 inhibit T cell activation by binding to the co-suppressor molecule CTLA-4; however, they promote T cell activation by binding to the corresponding co-stimulatory molecule CD28. The expression difference between these two pathways determines their dissimilarity. CD80/CD86 is mainly expressed in APCs, while the expression of CD155 is very extensive. Although not expressed in DCs, the expression of CD155 in various non-professional APCs, such as vascular endothelial cells, fibroblasts, and tumour cells, has been detected (32, 33). When autoimmune diseases occur, the tissue that is infiltrated by T cells contains mainly non-professional APCs, suggesting that the mechanism underlying the function of the TIGIT-CD226-CD155-CD112 network in immunity needs to be further studied(34).

We previously evaluated the therapeutic role of TIGIT-Fc in a model of murine lupus. However, in a previous study, the TIGIT-Fc protein we used was a fusion protein containing the murine TIGIT-ECD linked to the murine IgG2a chain without any modification of the Fc domain. In this study, we used a LALA-PG Fc variant to eliminate potential cytotoxicity. This is important because tDCs, which are key players in maternal immune tolerance, express high levels of the TIGIT functional receptor CD155. Moreover, a TIGIT-Fc with a wild-type Fc domain shows a strong effect on ADCC and C1q binding activity in vitro, further verifying the need to re-engineer the molecular structure of the TIGIT therapeutic protein.

We found that there was no significant difference in TIGIT expression between the decidual and peripheral lymphocyte cell subsets, as reported previously(1). Since CD155 is a receptor extensively expressed on human trophoblasts and decidual cells, it is not surprising that TIGIT participates in the development of normal human early pregnancy. The extravillous trophoblasts are in close contact with resident DCs in the decidua, usually in the decidua basalis(35). Moreover, our data show that high levels of secreted IL-10 were observed when decidual DCs were treated with TIGIT-Fc, endowing decidual DCs with the ability to induce the total decidual CD4^+^ T cells to produce increased levels of IL-5, IL-4, and IL-10 and minimal inflammatory cytokines, such as TNF-α and IFN-γ. However, TIGIT-Fc alone did not induce peripheral immature MDDCs to produce a high level of IL-10, suggesting differences in the cellular statuses of tDCs and immature MDDCs. Moreover, we further showed that treatment with TIGIT-Fc in the embryo resorption model (DBA/2J-mated CBA/J female) restored tolerance through the expansion of Foxp3^+^ Tregs and prevented foetal loss. Tregs are critical for normal pregnancy by regulating fetomaternal tolerance. Studies have demonstrated the prevention of foetal loss by the adoptive transfer of Tregs in an embryo resorption model and the high levels of embryo resorption induced by the depletion of Tregs in an allogeneic pregnancy model(36, 37). Foxp3^+^ Tregs are generated from naive CD4^+^ T cells in the periphery and thymus (38). Recently, extrathymic Foxp3^+^ Tregs have been reported to play an essential role in fetomaternal tolerance, while foetal-specific Tregs expanded rapidly and further induced tolerance during the subsequent pregnancy (38, 39).

Interestingly, our data clearly proved that progesterone upregulates TIGIT expression at the transcriptional level in decidual CD4^+^ and dNK cells in both humans and mice. Furthermore, we discussed the mechanism of the upregulation by progesterone of TIGIT gene expression by examining whether the identified putative PRE motifs participated in mediating the effect of progesterone/progesterone receptors on TIGIT promoter activity. The site-directed deletion mutagenesis study indicated that a mutation in the middle of the four putative PRE sites in both the human and mouse gene promoters resulted in the notable reversal of the effect of progesterone on TIGIT-luc activity but did not lead to an effect on basal TIGIT-Luc activity in the absence of progesterone. We also detected an interaction between progesterone and the TIGIT promoter at the molecular level.

Overall, the immunoregulatory role of TIGIT-Fc may provide pivotal insights into the regulation mechanisms of maternal immunity that allow successful pregnancies. Moreover, our data on the immunoregulatory therapeutic efficiency of TIGIT are important for understanding the function of fusion protein treatment under pathogenic conditions. TIGIT-Fc-based bio-therapy is expected to be a potent approach for the treatment of recurrent miscarriage with an immune aetiology.

## Acknowledgments

This study was financially supported by the National Natural Science Foundation of China (grant no. 81773261 and 81602690); Shanghai Rising-Star Program (19QA1411400); Shanghai Sailing Program (19YF1438600); Shanghai Chenguang Program (17CG35); Military Medicine Special Grant of Second Military Medical University (grant no.2017JS01); a General Financial Grant from the China Postdoctoral Science Foundation (grant no. 2016M593006), and postdoctoral scientific research funds of Second Military Medical University.

