## Supplementary for "TIGIT-Fc Promote Immune Tolerance at the Feto-maternal Interface"

**Supplementary Information for:**

***Tightening Maternal Rejection via TIGIT-Fc***

Contents	Page

Supplementary Methods	2

Supplementary Figures	3

Figure S1. Binding of Fusion proteins to human antigens.	3

Supplementary Tables	4

Table S1. Selected analytical data and pharmacokinetic parameters of recombinant fusion proteins in mice.	4

Supplementary Table 2. Primer sequences	5

### Supplementary Methods

**Real-time polymerase chain reaction (PCR)**

Total RNA was isolated with the RNeasy Mini Kit (Qiagen) according to the manufacturer’s instructions. Real-time quantitative PCR was performed with an ABI PRISM 7900HT instrument with the commercially available TaqMan probes Hs00545087 and Mm03807522. Data were normalized to β-actin, which served as an endogenous control, and analysed using SDS v2.3 (Applied Biosystems).

**IgG Biological Effect Assays**

For the in vitro ADCC assay, SupT1 cells expressing murine PVR and A431 cells (high-expressing PVR cells) were labelled with 5 mM carboxyfluorescein succinimidyl ester (CellTrace CFSE Cell Proliferation Kit, Life Technologies) and co-cultured with murine or human macrophages overnight, respectively, at the indicated ratios in the presence of TIGIT-Fc fusion proteins.

For C1q ELISA, an ELISA sandwich-type immunoassay was used to analyse the binding of the different fusion proteins to C1q. Each fusion protein was coupled to a hydrophobic Maxisorp 96-well plate at eight different concentrations between 10 and 50 μg/ml. After washing, the C1q samples were incubated on the plate to allow C1q to bind to the fusion proteins. The bound C1q molecules were further washed and detected by anti-C1q antibodies followed by an HRP-labelled secondary antibody.

### Supplementary Figures

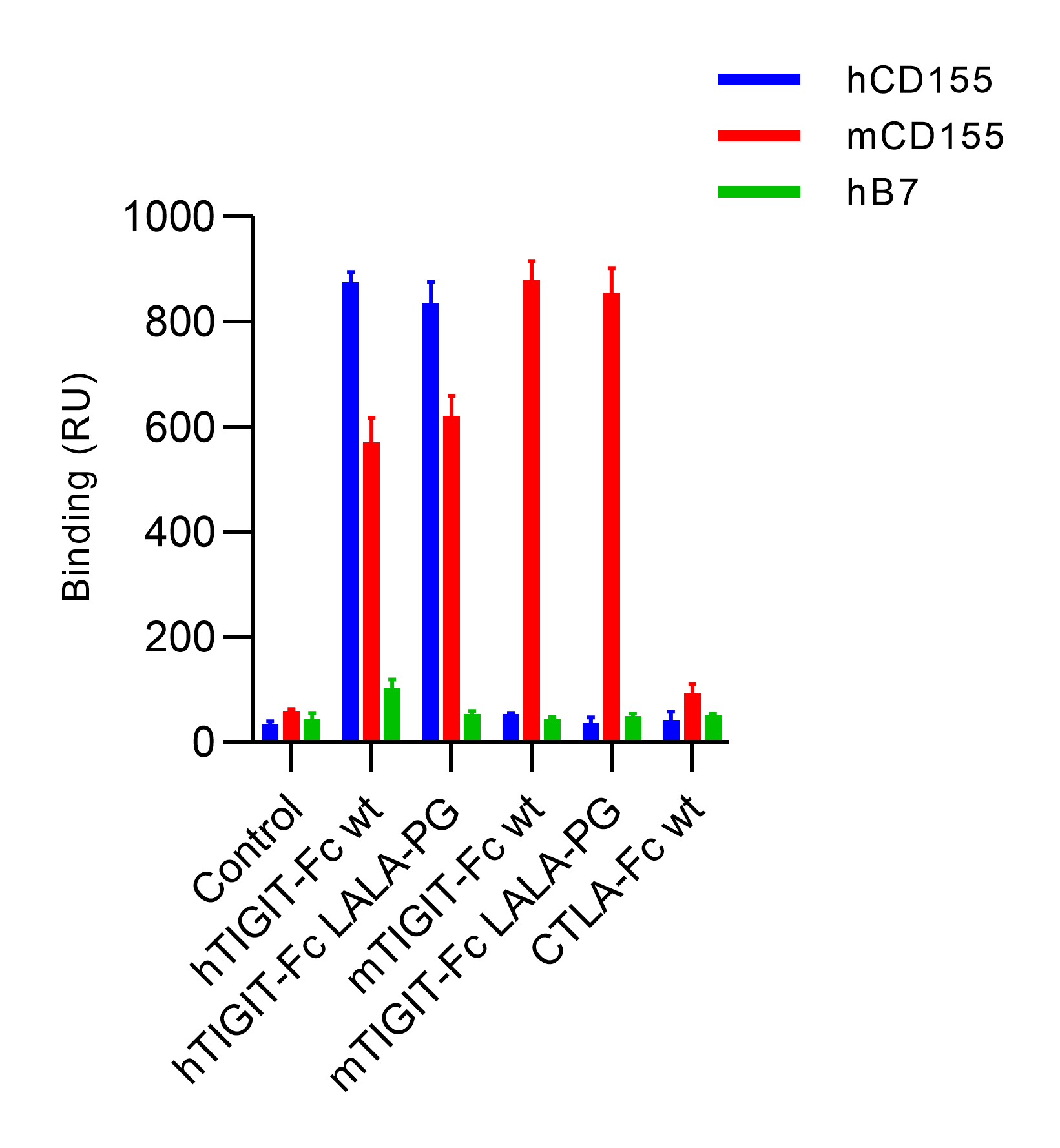

#### **Figure S1. Binding of Fusion proteins to human antigens.**

Fusion proteins was tested for binding to immobilized hCD155, mCD155 or B7 protein using surface plasmon resonance on a BIAcore 2000 instrument. Binding was quantified as an increase in RU at 60 s after the end of injection compared with a baseline established 20 s before injection.

### Supplementary Tables

#### Table S1. Selected analytical data and pharmacokinetic parameters of recombinant fusion proteins in mice.

| Parameter^a^ | CTLA4-Fc wt | hTIGHT-Fc wt | hTIGIT-Fc LALA-PG | mTIGIT-Fc wt | mTIGIT-Fc LALA-PG |
| --- | --- | --- | --- | --- | --- |
| HMW formation after storage(% SEC area)^b^ | < 0.1 | < 0.1 | < 0.1 | < 0.1 | < 0.1 |
| LMW formation after storage(% SEC area)^b^ | < 0.1 | < 0.1 | < 0.1 | < 0.1 | < 0.1 |
| AUC (day μg ml^-1^) | 492.20 | 524.63 | 489.03 | 517.63 | 495.33 |
| *T_1/2_(*day) | 5.82 | 6.66 | 4.90 | 4.95 | 5.91 |
| CL (ml day^-1^ kg^-1^) | 7.31 | 6.98 | 7.97 | 7.72 | 7.68 |
| VSS(ml kg^-1^) | 81.14 | 77.43 | 77.35 | 70.83 | 78.61 |

^a^ Pharmacokinetic parameters were calculated using a noncompartmental analysis. AUC, area under the concentration versus time curve; t1/2, half-life; CL, clearance; VSS, steady-state volume of distribution.

^b^ Quiescent storage for 3 wk, 40 °C, 1 mg/mL

#### Supplementary Table 2. Primer sequences

| Sets | Forward | Reverse |
| --- | --- | --- |
| Human Set1 | 5’- CACAAGTGGCTTTGGAACTC -3’ | 5’- ATGTTAGGAGAGAAGGGCG -3’ |
| Human Set2 | 5’-CAGATCACGGGCAGTCCTTT-3’ | 5’-AGCTGTCTCACAGCCTTAGC-3’ |
| Murine Set1 | 5’- ATCTCTCTGGGCAAGAAGTC -3’ | 5’- CTGGTGGGAAGAACACAAC -3’ |
| Murine Set1 | 5’-GTGCGTCACCTACCCTTGGAA-3’ | 5’-AAAGCCACATAGGGTGAGCC-3’ |
